# Beyond tree-like evolution in ostariophysan fishes

**DOI:** 10.64898/2026.11.09.765406

**Authors:** Gustavo A. Ballen, Claudia Solís-Lemus, George P. Tiley, Claudio Oliveira

**Affiliations:** Instituto de Biociências de Botucatu, Universidade Estadual Paulista “Júlio de Mesquita Filho”, Botucatu, SP, Brazil; Wisconsin Institute for Discovery, University of Wisconsin, Madison, WI, United States; Department of Plant and Microbial Biology, North Carolina State University, Raleigh, NC, United States

## Abstract

Phylogenetic trees have long been a powerful model of the evolutionary history. However, key processes in speciation, such as introgression and hybridization are difficult to depict with strictly binary trees alone, even with modern discordance metrics and post-hoc tests. Phylogenetic networks offer an intuitive way to estimate and interpret these reticulate events, as conflicting signal from gene trees should become evident as hybrid edges. Phylogenetic networks raise a number of challenges over tree-based analyses though, including the estimation of uncertainty and evaluating confidence among multiple competing network hypotheses. Here, we develop a new approach for assessing branch and topological support on networks using composite likelihood, and explore the utility of such statistics by revisiting competing evolutionary hypotheses in ostariophysan fishes. Our results suggest that a tree-like topology is not the best model for ostariophysans. A phylogenetic network with four reticulation events involving early-diverging lineages was the best alternative scenario. Our branch support metrics demonstrated non-negligible levels of uncertainty in the placement of those reticulation events though. An important question though, is if the uncertainty in the placement of these reticulation edges affects the qualitative conclusions of organismal evolution. To this end, we apply our network hypotheses to the biogeographic history of ostariophysans. Since no biogeographic models for networks exist, we implemented a parsimony approach using Fitch’s algorithm for ancestral character state reconstruction. This represents the first biogeographic reconstruction for networks, and while simplistic, is a basis for further development. Biogeographic results are generally consistent with previous research on the presence of ancestral lineages in Laurasia and Gondwana post the break-up of Pangea, but also identify some cases of long-distance dispersal.

## 2 Introduction

Ostariophysans have long been a group of fishes with an uncertain evolutionary history (Figure 1; Oliveira, 2025). Although multiple independent investigations support monophyly of the ostariophysan superorder (Fink & Fink, 1981; Orti & Meyer, 1996; Saitoh et al., 2003; Chen et al., 2013; Arcila et al., 2017; Betancur-R et al., 2017; Betancur-R. et al., 2019; Chakrabarty et al., 2017; Mirande, 2017; Dai et al., 2018; Hughes et al., 2018; Simion et al., 2020; Lenglin et al., 2025; Liu et al., 2025; Brownstein et al., 2026; Hays et al., 2026), the monophyly of some of the orders (e.g., Characiformes) and the relationships among orders have been controversial using both morphological and molecular datasets.

**Figure 1:**
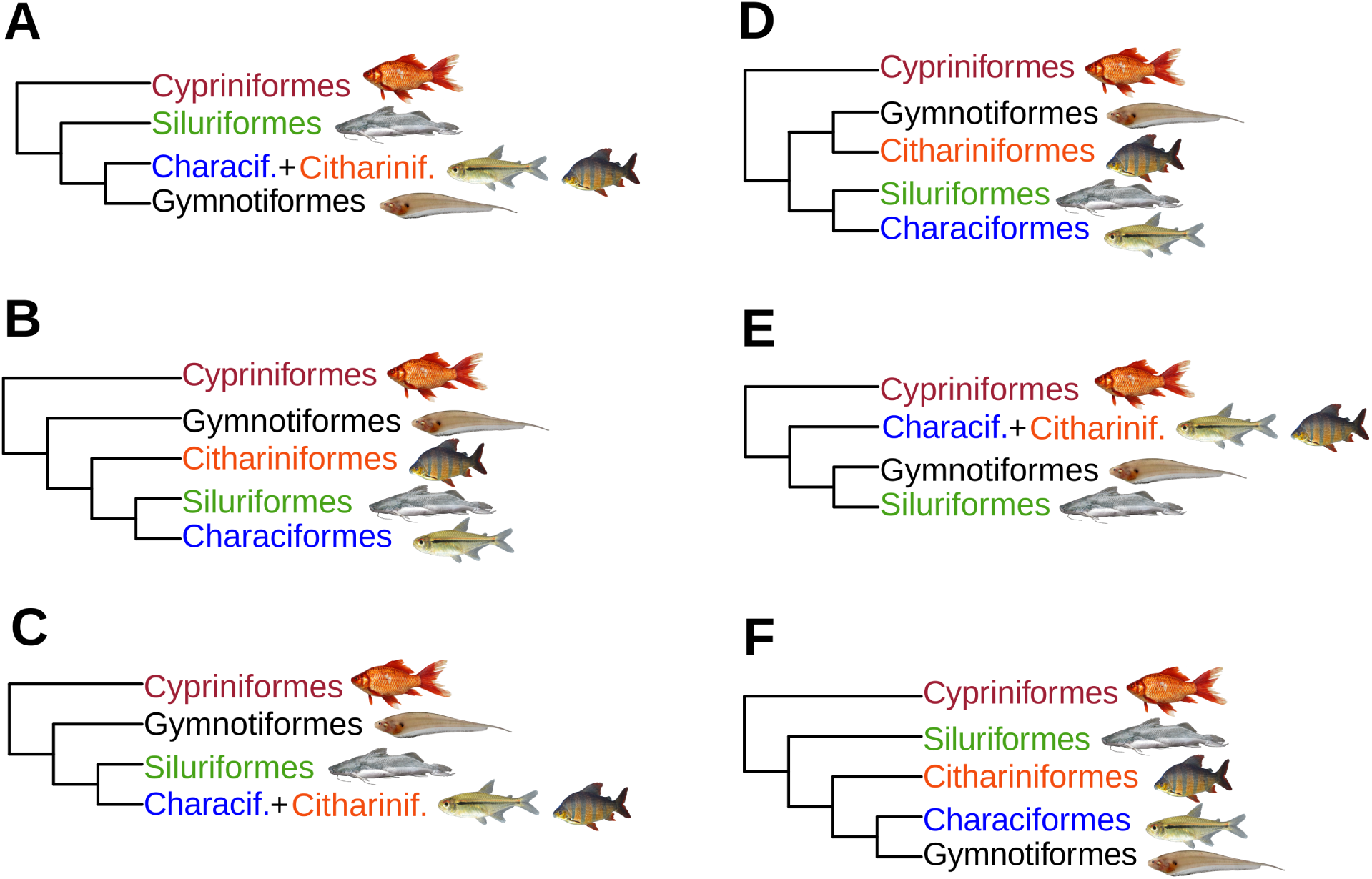
Phylogenetic relationships among ostariophysan orders have been found to be elusive in previous studies. Here we consider Characiformes as distinct from Cithariniformes. A) Characiformes+Cithariniformes sister to Gymotiformes (Saitoh et al., 2003). B) Characiformes and Cithariniformes non-monophyletic, Characiformes sister to Siluriformes (Chen et al., 2013; Chakrabarty et al., 2017; Mirande, 2017; Simion et al., 2020; Brownstein et al., 2026; Hays et al., 2026). C) Characiformes+Cithariniformes sister to Siluriformes (Lenglin et al., 2025). D) Characiformes and Cithariniformes non-monophyletic, Characiformes sister to Siluriformes and Cithariniformes sister to Gymnotiformes (Liu et al., 2025). E) Characiformes+Cithariniformes sister to Siluriformes+Gymnotiformes (Fink & Fink, 1981; Arcila et al., 2017; Betancur-R et al., 2017; Hughes et al., 2018; Betancur-R. et al., 2019). F) Characiformes and Cithariniformes non-monophyletic, Characiformes sister to Gymnotiformes (Orti & Meyer, 1996; Dai et al., 2018).

Fink & Fink (1981) proposed the historical relationships among ostariophysan orders, which are always recovered using morphological datasets and sometimes using phylogenomic data (Arcila et al., 2017; Hughes et al., 2018). Betancur-R et al. (2017) updated previous classifications of Bony Fishes using genomic-scale sampling for thousands of species. They recovered a ladderized ostariophysan tree where Gymnotiformes is sister to Siluriformes, and then the orders Characiformes, Cypriniformes, and Gonorynchiformes at the base of the subtree.

Chakrabarty et al. (2017) addressed the relationships of ostariophysans by using 567 Ultraconserved Elements (UCEs) as well as combined analysis of sequence and morphological data. The order Characiformes was recovered as non-monophyletic, but morphology favored the monophyletic hypothesis given the data of Fink & Fink (1981). Otherwise, the UCE results of Chakrabarty et al. (2017) recovered the same order-level relationships found earlier by an earlier study using five Sanger loci (Chen et al., 2013). Despite the apparent consistency across molecular phylogenetic investigations, subsequent phylogenomic studies have also found high levels of discordance between gene trees and species trees. Specifically, the orders Characiformes, Gymnotiformes, and Siluriformes have been found to have strong topological conflict in their early-diverging sections. Alda et al. (2019) carried out an analysis of gene-tree to species-tree discordance in the order Gymnotiformes using UCE sampling and found that different filtering strategies resulted in general agreement, although it pinpointed areas of the gymnotiform tree where discordance would lead to topological uncertainty. They found that concatenated and coalescent-based analyses revealed high gene-tree discordance. Their results highlight the importance of using coalescent-based methods when the presence of discordance is rampant, although by that time the methods for phylogenetic network inference were not capable of processing large taxon sample size. In summary, their results point to phylogenetic network inference as a promising tool for analysis of UCE sampling under gene-tree to species-tree conflict. Betancur-R. et al. (2019) assessed the monophyly of Characiformes using phylogenomic sampling composed by UCEs and exon capture from previous studies and tested hypotheses of characiform monophyly using gene genealogy interrogation (Arcila et al., 2017). Rivera-Rivera & Montoya-Burgos (2018) explored the sources of discordance among basal interrelationships in the order Siluriformes using ten loci and suggested that rate heterogeneity may explain the conflicting position of the Loricarioidei as the most basal suborder in contrast to previous studies which have found the Diplomystoidei to be the earliest branch among living representatives. This approach however does not explore the possibility that gene–tree to species–tree discordance can be caused by incomplete lineage sorting at a genomic scale, or whether reticulations can explain these conflicting topologies. Arroyave et al. (2013) focused on the interrelationships of the characiform suborder Citharinoidei and their timetree including ostariophysan outgroups showed that siluriforms and characiforms form a clade diverging since the early Cretaceous using a concatenated dataset of seven loci.

Arcila et al. (2017) proposed gene genealogy interrogation as a practical method for addressing topological conflict that stems from gene tree estimation error. They used primarily the otophysans (ostariophysans excluding Gonorynchiformes) as case study as well as other groups to assess the potential of the method. They recovered the same interrelationships as Betancur-R. et al. (2019), reinforcing a tree-like view of the evolutionary relationships among Otophysans. They suggest that either incomplete lineage sorting or the anomaly zone are not a source of conflict but rather gene tree estimation error. They recovered strong evidence for the monophyly of the orders as well as for the traditional classification of the group. However, they still recover several clades where topological conflicts cannot be resolved even when picking an optimal tree using topological testing, which suffers from the drawbacks of frequentist approaches such as inflating type I errors when carrying out multiple tests. Thus, even when the overall relationships might be robust, several orders have been reported to persist showing conflicting interrelationships (e.g., characiform and siluriform suborders, and major clades of the Gymnotiformes, Arroyave et al., 2013; Alda et al., 2019; Arcila et al., 2017). In these instances, phylogenetic networks can be a powerful tool for better understanding the reasons for topological uncertainty if incomplete lineage alone is not sufficient for explaining gene tree variation.

The non-branching processes of hybridization/introgression have long been recognized as important evolutionary phenomena that produce non-bifurcating events. This can manifest in species trees as poorly supported nodes, which do not necessarily represent noisy data but rather a deviation from the tree model in favor of a network (Solís-Lemus & Ané, 2016; Jiao et al., 2020). Although phylogenetic network inference has seldom been used for estimating interrelationships of fishes, the role of introgression and hybridization has already been suggested as important in explaining topological conflict in phylogenetics and genomics (Bangs et al., 2018; Meier et al., 2023; Meyer et al., 2017). Here, we anticipated that disagreements in ostariophysan evolution were due, at least in part, to historical gene flow among common ancestors. If so, phylogenetic networks should be a better depiction of the evolutionary relationships among ostariophysans. We generated new UCE data to estimate networks for ostariophysans, and implemented new bootstrapping approaches for evaluating uncertainty in networks too. Network estimates were used to evaluate potential associations between reticulate evolution and biogeography under a parsimony model.

## 3 Materials and methods

### 3.1 Taxon sampling

A comprehensive sample of ostariophysans and outgroup related taxa was compiled for a total of 231 terminals in the dataset. The outgroup includes the orders Alepocephaliformes (7 spp.), Argentiniformes (2), Clupeiformes (15), Notachanthiformes (1), Anguilliformes (1), Elopiformes (1), Osteoglossiformes (2), and Amiiformes (1). The ingroup includes representatives of all ostariophysan orders, recognizing Characiformes and Cithariniformes as separate. Our taxon sampling comprises the orders Characiformes (60 spp.), Cithariniformes (12), Cypriniformes (35), Gonorynchiformes (11), Gymnotiformes (13), and Siluriformes (71). We included at least one representative per family in each order, often including more than one per genus when available, or a single sample in monogeneric families (e.g., Tarumaniidae). Sequences were generated by our research group at UNESP Botucatu or retrieved from GenBank. Details are available in the Supplementary Table S1.

### 3.2 Phylogenomics

#### 3.2.1 DNA Extraction and UCEs Sequencing

Total DNA was extracted from muscle samples preserved in ethanol with the DNeasy Tissue Kit (Qiagen) following the manufacturer’s instructions. The quantification was performed using the Qubit dsDNA BR Assay Kit (Invitrogen–Life Technologies). We used sets of probes developed for ostariophysan fishes (Faircloth et al., 2020) which generated sequence data with 2700 UCE loci. DNA libraries were prepared following the protocol modified from Nextera (Epicenter Biotechnologies). The enrichment steps were performed using the solution-based target method of Faircloth et al. (2012). The number of PCR cycles after the fragmentation reaction was increased to 20 (Faircloth et al., 2013). We used the Nextera library preparation protocol for in vitro transposition followed by PCR to suppress DNA and attach sequence adapters following Adey et al. (2010). We followed the transposase-mediated library prep protocol of the Epicenter Nextera kit that produced libraries that insert sizes with an average of 100 bp (95% CI: 45 bp Adey et al., 2010). The samples were initially sheared with a Qsonica Q800R sonicator during library preparation and then selected for a modal length of approximately 500nt using a dual-step SPRI bead cleanup. We converted the DNA into Illumina sequencing libraries with a slightly modified version of the NEBNext Ultra (DNA Library Prep Kit for Illumina). The libraries were amplified using KAPA HiFi HotStart ReadyMix (Kapa Biosystems) for 6 cycles using the thermal profile recommended by the manufacturer and double P7 and P5 primer indexes (Kircher et al., 2012). After purification with SPRI beads, the libraries were quantified with the nt-iT (Picogreen dsDNA Assay kit (ThermoFisher) assay kit. The enriched pools were composed of 100 ng from each of the 8 libraries (800 ng of total input material) using the MYbaits Target enrichment system (MYcroarray) following manual version 3.0. After cleaning (i.e., the bead-bound library), it was resuspended with the recommended solution and amplification for 10 cycles using the P5 / P7 primer pair and KAPA HiFi. After purification, each captured library pool was quantified with PicoGreen and it was combined with all other pools in an equimolar ratio before sequencing. The sequencing was performed using HiSeq paired-end 100 bp using v4 chemistry.

#### 3.2.2 UCE sequence analysis

The details of the UCE sequence analysis are available in the PHYLUCE documentation (Faircloth, 2016). After sequencing, adapter contamination, low quality bases, and sequences containing ambiguous bases were cut using the software illumiprocessor (Faircloth, 2011). We assembled the contigs using SPAdes (Prjibelski et al., 2020). After assembly, we used the program match contigs to probes.py, present in the PHYLUCE software package (Faircloth, 2016), integrating LASTZ (Harris, 2007) to align species-specific contigs to the UCE probe set. This last program creates a relational database of matches between UCE loci for each taxon. We used get match counts.py to search the database and generate a separate file for the UCE loci that identifies all the taxa. Then, we aligned multiple sequences using MAFFT (Katoh & Standley, 2013). Because of the computational burden of processing all the samples at once with PHYLUCE, we captured UCEs in blocks of terminals, often around 30 species, and then combined the alignments using yafastac (Ballen, 2025), and then we filtered the UCE alignments which contain 70% or more taxa, which recovered 1150 loci.

### 3.3 Phylogenetic network inference

#### 3.3.1 Subsampling approach

Phylogenetic network inference is very challenging as the parameter space grows massively with the number of terminals (Kong et al., 2025). Current methods tend to scale poorly as the number of terminals increases, therefore restricting inference of networks to datasets of a handful of species. We integrate several novel methods (Figure 2) that help improve computational efficiency while scaling to the number of taxa, which is large for current standards in phylogenetic network inference. First, gene trees were inferred using IQTREE 3 (Wong et al., 2025). An initial species tree was estimated using quartets with TREE-QMC (Han & Molloy, 2023), and then rooted using phyx (Brown et al., 2017). Then, a tree of blobs was estimated using TREE-QMC (Dai et al., 2025) in two steps: First, an exhaustive search for p-value was carried out on each branch where the quartet and star tree hypotheses were tested, recording the number of observed quartets, their proportion, and p-value using TINNIK (Allman et al., 2024) inside TREE-QMC. Then, we chose thresholds for collapsing branches using first the total number of gene trees where the branch was present, then the proportion between the two least frequent quartets, and finally the p-value. Second, those branches flagged with the thresholds were collapsed into polytomies representing blobs. Given the strategy above, we set the hyperparameters *α* and *β* to 1e-11 and 0.95 respectively, because these retained branches with a large number of observed gene trees, and non-trivial proportion between the two least frequent qCFs.

**Figure 2:**
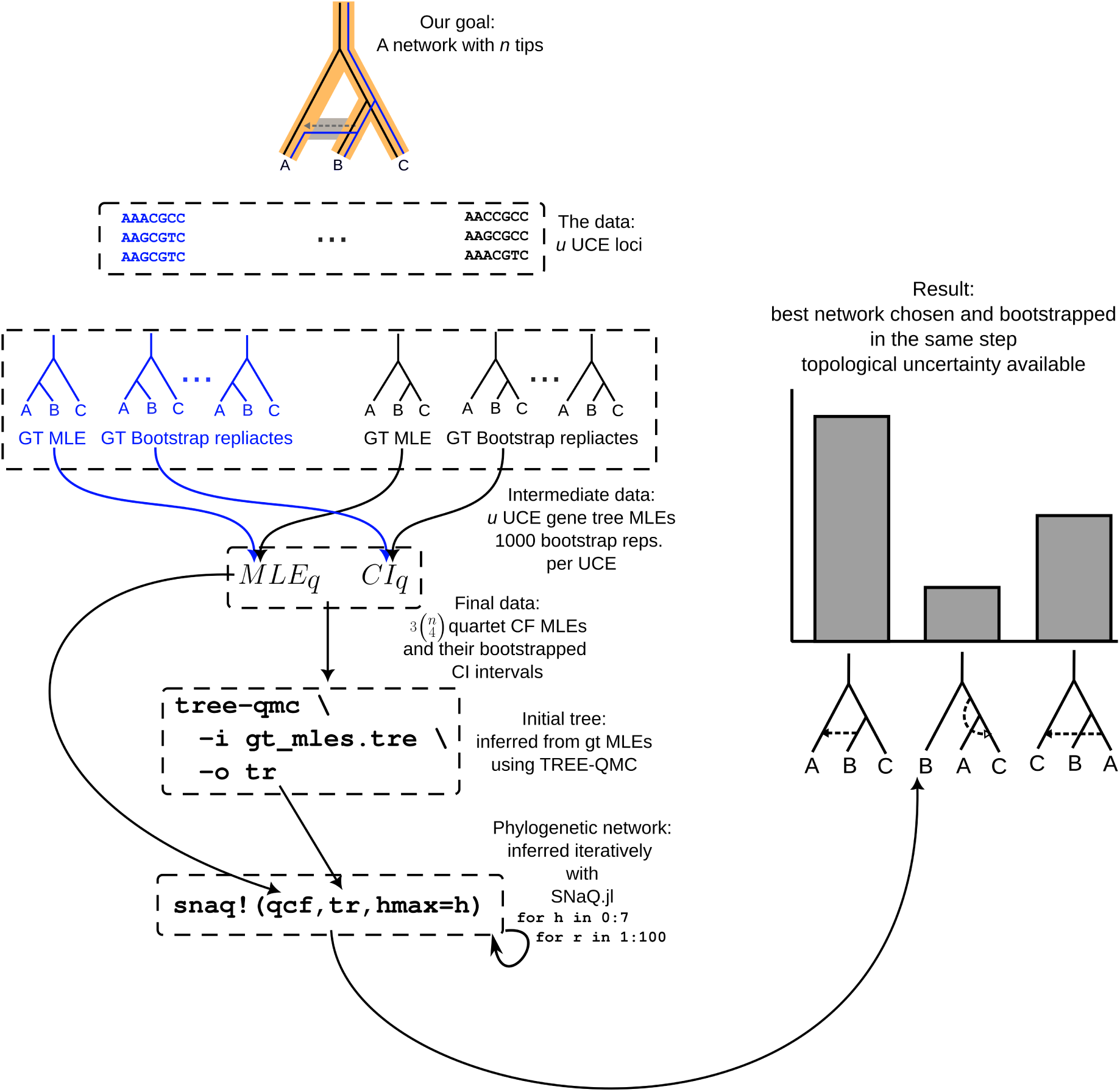
Workflow proposed in this work. We start with *u* ultraconserved element loci (UCE) (1150 in this study), and then estimate a maximum likelihood (MLE) and 1000 bootstrap replicates for each *u* gene tree (GT). Here two gene trees from the phylogenetic network under the multispecies coalescent with introgression are depicted for illustrative purposes, one evolving through the major tree (black) and one through the reticulation (blue). Both MLEs and bootstrapped GTs are used for estimating MLE quartet (q) concordance factors (CF) as well as the 95% confidence interval (CI) for each qCF. Then an initial tree is estimated using the GT MLEs using TREE-QMC. Iterative runs using SNaQ are carried out with maximum number of hybrid edges from 0 to 7, with 1000 independent runs for each of these. The multiple runs are then summarized for estimating topological uncertainty using the function topoFreq in the tbea package, choosing the best network, and summarizing bootstrap support values on the focal network.

The tree of blobs inferred showed that possible signal for reticulate evolution is restricted to an internal section of the tree that involves the bases of the orders Characiformes, Cithariniformes, and Siluriformes. We then subsampled the gene trees so that only portions which clear sign of reticulation were preserved. Care was taken to preserve the distinction between internal and external branches by replacing a resolved subtree with a couple taxa that represent the same node. Also, the outgroup was pruned to a single representative of Cypriniformes because the tree of blobs showed that neither Gymnotiformes nor Cypriniformes were involved in possible reticulations, and therefore a single representative of each was preserved. This produced a smaller problem of 43 terminals. Gene trees from the subsection above were subsampled only to include the list of 43 terminals. An initial tree was estimated using quartets with TREE-QMC. Concordance factors were calculated using PhyloNetworks.jl.

#### 3.3.2 Phylogenetic network inference incorporating data uncertainty

Introducing uncertainty in data for phylogenetic network inference is often restricted to the generation of bootstrap replicates (Solís-Lemus et al., 2017). In these applications, confidence intervals are constructed for the quartet concordance factors and then SNaQ samples a given value per bootstrap replicate, carry out network inference, and then summarize the results on a focal network, often the one selected with score heuristics (Tiley et al., 2024). Here we propose a different approach at inference itself (Figure 2). Instead of using a fixed collection of quartet concordance factors, one can run multiple independent analyses, one with values sampled from the concordance factors, and then summarize the collection of best networks inferred this way. Afterwards, summarization on a focal topology is a trivial matter as the multiple inferred topologies are already available. It is similar in spirit to sampling from a posterior distribution of topologies in a Bayesian analysis, and then summarizing the same distribution on a focal topology, for instance using the highest posterior probability tree (Bouckaert et al., 2019).

So far SNaQ allows two different approaches when incorporating uncertainty: 1) to use the TICR pipeline for calculating confidence intervals on quartet concordance factors from posterior gene tree samples, or 2) to use a vector of bootstrapped gene trees. The former approach forces one to use Bayesian inference for gene tree collections, while the latter forces the user to have the same set of terminals in all bootstrapped trees. In the latter case, gene trees with incomplete sampling, which are quite frequent in phylogenomic datasets, force the bootstrapped network to have the set of terminals common to all trees, and therefore can potentially lose multiple terminals. We have implemented a method for calculating the confidence intervals of quartet concordance factors from arbitrary collections of replicates for gene trees, for instance ultrafast bootstrap ones estimated with IQTREE, so that one is not forced to use any approach or program, be it Bayesian or maximum likelihood. The function confintqCF genetrees uses a vector of point estimate gene trees (of type Vector*{*HybridNetwork*}*), and a vector of replicaces per gene trees (of type Vector*{*Vector*{*HybridNetwork*}}*), and returns a concordance factor table with columns for the confidence intervals and their point estimates. The function is already available in SNaQ.jl v1.2.0 onwards (Kolbow et al., 2025).

The method above poses a number of challenges to standard point estimation as done in SNaQ. First, we no longer have a single best network but rather a collection of best networks. Second, we have in one shot network inference and uncertainty assessment. Therefore, we need a different way to pick the best network. We propose examining the topological frequency to pick the focal network to visualize and to map bootstrap replicates onto, which gives us also access to topological uncertainty in general. A similar approach is available for Bayesian tree inference as implemented in tbea (Ballen & Reinales, 2025). The function topoFreq from the tbea package takes a vector of topologies and identifies clusters of identical topology. Alternatively, one can use an arbitrary pairwise distance matrix to find these clusters (available from tbea v1.8.0). In this way we can find the most frequent ones, or any set of cumulative frequency, say the set of 95% topologies. Because the method was originally implemented with tree topologies in mind, it uses the Robinson-Foulds distance. However, one can also calculate a pairwise topology distance matrix using the *µ*-distance (Ribas et al., 2026) which can be used for non-level-1 networks, or the hardwired cluster distance for level-1 networks (Maxfield et al., 2025), allowing the use of the topoFreq function for phylogenetic networks too. In order to identify clusters for which we can trace hybrid edges, that is, for which the major tree remains the same, we first clustered together networks that share the same major tree and then estimated topological uncertainty on such sets. The rationale behind this is that methods based on the multispecies coalescent process (e.g., ASTER-Astral, TREEQMC, SNaQ; Han & Molloy, 2023; Zhang et al., 2025; Li & Ané, 2026) should have already identified the tree-like part of the topology, and therefore further accepted topological moves that are not focused on hybrid edges are probably errors rather than true refinements during network inference. When the tree-like part of the topology is inconsistent across networks, hybrid edges may be impossible to trace between networks, and thus such bootstrap replicates become useless.

Because the largest empirical problem attempted with SNaQ v1.0 and previous versions was around 30 terminals, and ours is larger (43 terminals), we used the newest version of SNaQ *>*v1.1.1 which is considerably faster, with three techniques for accelerating analyses either alone or in combination: 1) using 10 threads per run, so that the composite likelihood is parallelised over the available threads; 2) using subsampling of quartets set to 50% and then to 10%; and 3) removing uninformative quartets close to equal proportions of 1/3. We then compared both runtime and the resulting runtimes and composite likelihoods and chose the best scheme for submitting 1000 independent runs over high throughput computing on the Open Science Grid which were then used for selecting the best composite likelihood. We ran subsequent calls to the function snaq! using hmax = 0 to 7 with 200 independent runs, and chose the best network with the heuristic saturation rule of the composite likelihood profile (Tiley et al., 2024). Here, we applied the suggested modification for incorporating data uncertainty and measuring topological variation among analyses.

### 3.4 Sources of phylogenetic incongruence

We estimated a species tree using ASTRAL-4 (Tabatabaee et al., 2023), part of the ASTER suite (Zhang et al., 2025), using the annotation argument -u 2. Then, we used phytop (Shang et al., 2025) for quantifying topological conflict from different sources. The quartet frequencies for each branch and the proportion of gene trees that can be explained either by incomplete lineage sorting (ILS-e) or hybridization/introgression (IH-e) were used for visualization of gene tree variation. The species tree estimated with ASTRAL-4 was then compared with the major tree implied by the selected phylogenetic network using a cotangleplot from phytools (Revell, 2024).

### 3.5 Biogeography on phylogenetic networks

Historical biogeography is a field with a long and rich tradition in evolutionary biology (Landis et al., 2026). Many of the methods use phylogenetic trees either as their model for depicting historical area relationships, or by allowing Markovian models that describe area dynamics through time to evolve on a phylogenetic tree. The state-of-the-art models in probabilistic biogeography fall within this second group, and require a time-calibrated phylogeny (Warnock & Wright, 2020). Phylogenetic network estimation is a much younger field than tree estimation, and therefore the methods for obtaining time-calibrated networks are still in active development. Additionally, while phylogenetic comparative methods now have many software packages available for analyzing comparative data on trees, relatively limited options are available for neworks (Bastide et al., 2018).

The above limitation pose two challenges for biogeographic reconstructions. First, we need topologies where branch lengths represent absolute time and not coalescent units, which normally is achieved through relaxed-clock divergence time estimation not possible for our network. Second, because no biogeographic method available works on networks, probabilistic approaches would not be feasible even if we imposed divergence times from the major tree topology. However, the potential role of hybridization in biogeography has been suggested already in the literature (Waters et al., 2023), even in the absence of dedicated methods for examining such relationships. In order to overcome the lack of biogeographic methods working on phylogenetic networks, we used Fitch’s algorithm (Fitch, 1971) to estimate ancestral character states considering biogeographic areas as a multistate character. The underlying assumptions of Fitch’s algorithm are that transformations between character states are equally likely, and that the aim is to minimize the number of steps implied by the transformation of states on a topology. Fitch’s algorithm has been extended to phylogenetic networks (Kannan & Wheeler, 2012), and its implementation both for calculating the parsimony score of a character on a phylogenetic network and its ancestral states is already available in PhyloNetworks.jl (Solís-Lemus et al., 2017). Biogeographic areas occupied by the species in the present study are South America, Africa, India, North-East Asia, and South-East Asia.

## 4 Results

### 4.1 Tree of blobs

The tree of blobs (Figure 3) for the extended taxon sampling (231 terminals) recovered most of the ostariophysan phylogeny as tree-like, with five major non-tree-like and non-star-like branches (*p <* 10*^−^*^11^ and *p >* 0.95 respectively). Notable sections of the tree were the superfamily Pimelodoidea and Big Asia in the order Siluriformes (Sullivan et al., 2006), and the superfamily Anostomoidea, a clade comprising the families Serrasalmidae+Cynodontidae+Hemiodontidae, and the family Parodontidae of the order Characiformes. No major reticulate events were detected in the orders Gonorynchiformes, Cypriniformes, Cithariniformes, and Gymnotiformes. Importantly, the lack of monophyly in the Characiformes and Siluriformes suggested deep reticulation events towards the base of both orders.

**Figure 3:**
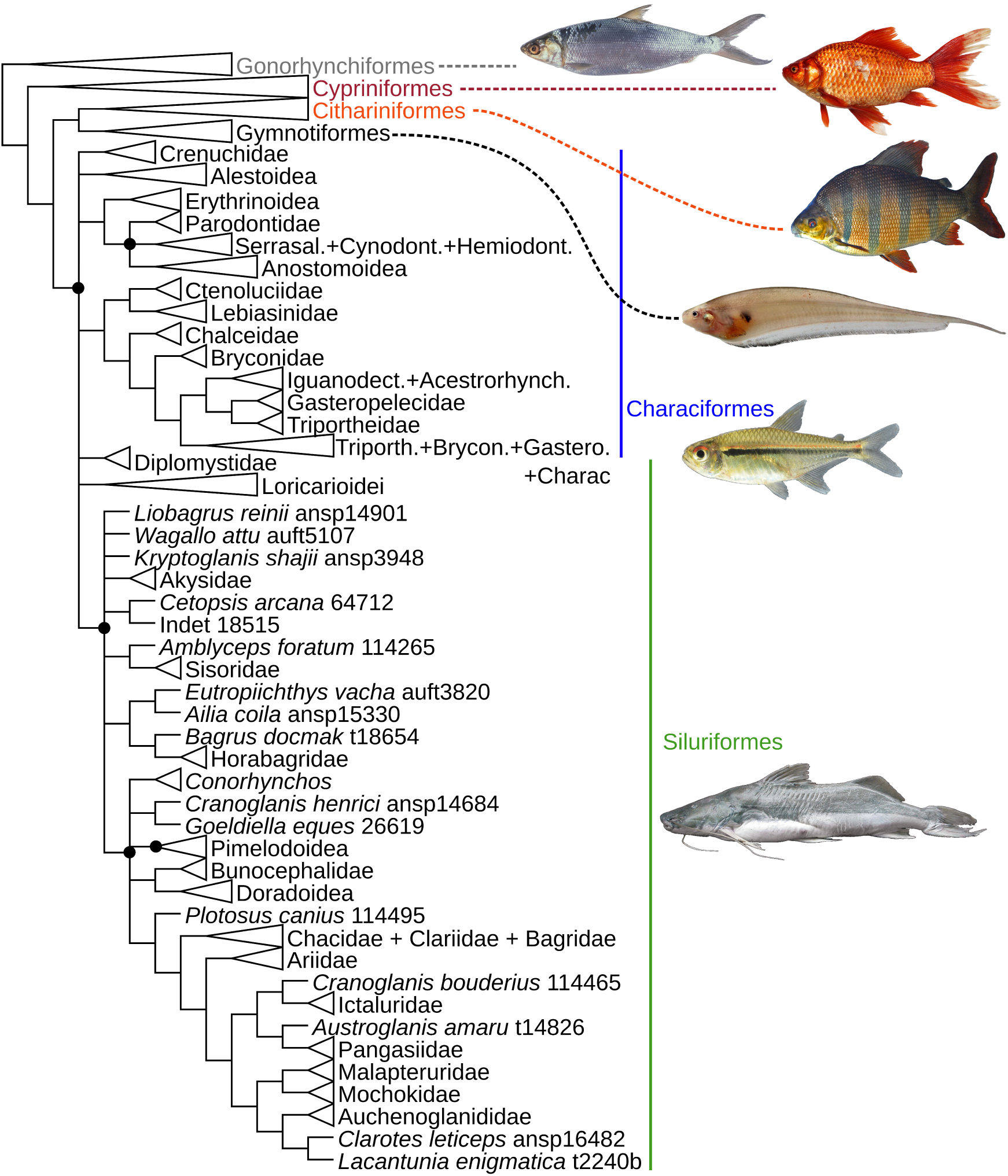
Tree of blobs with 231 terminal as inferred by the program Tree-QMC. Polytomies represent collapsed branches where *P* (tree-node) *<* 10*^−^*^11^ and *P* (star-node) *>* 0.95. These regions represent areas which are not tree-like and therefore show signal of reticulate evolution. Note that the monophyly of both Characiformes and Siluriformes is not recovered because of the extensive collapse of basal branches in both orders. Only relevant subtrees are shown where evidence of non-tree-likeness is present. Outgroup clades were removed. Abbreviated names in Characiformes are all families. Characiform superfamily composition follows Melo et al. (2022) (Erythrinoidea), Melo & Stiassny (2024) (Alestoidea), and Dillman et al. (2016) (Anostomoidea). Siluriform suborders and superfamilies follow Sullivan et al. (2006).

### 4.2 Phylogenetic networks

The best phylogenetic network (Figure 5A) for the reduced dataset with 43 terminals was found to show four reticulation events (Figure 4). Also, a strictly tree-like topology can be rejected for this dataset. Three of the reticulation events have unsampled ancestors, whereas one of them was a deep reticulation between sampled ancestral lineages. The inheritance probabilities (*γ*) for events was found to be between 2.2% and 6% (Figure 5A). Siluriform terminals were often the recipients of gene flow and deep portions of the topology were often donors, again pointing towards unsampled sources. A deep reticulation of interest originated from the suborder Loricarioidei into a clade comprising Asian siluriform genera defined by *Bagarius yarrelli* and *Akysis patrator*.

**Figure 4:**
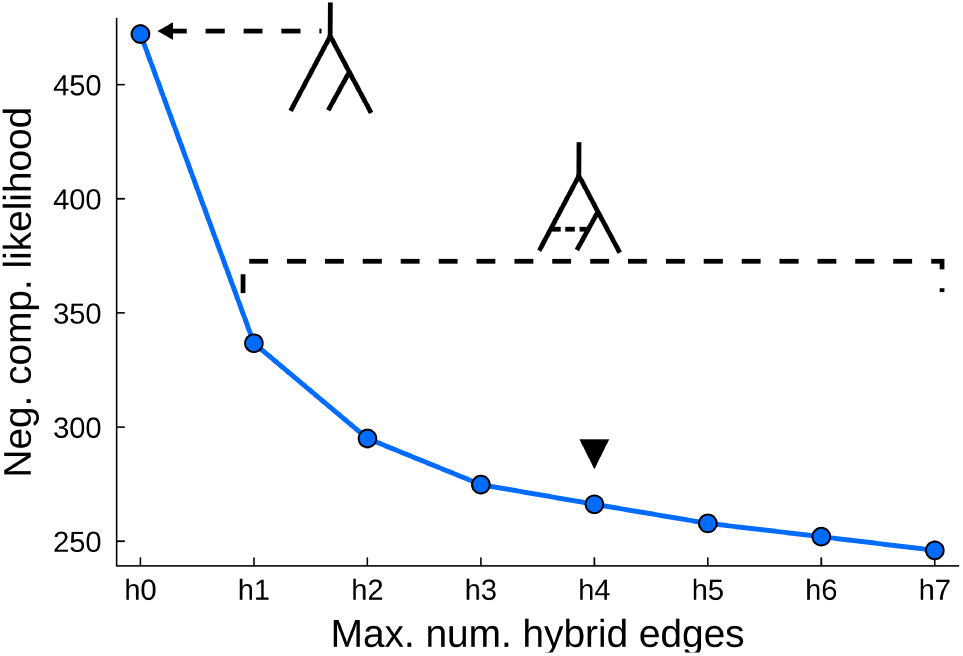
Profile of negative composite likelihood on the vertical axis as a function of maximum number of hybrid edges per analysis on the horizontal axis. Gradient stabilizes at four hybrid edges (*h* = 4, downward triangle) which is chosen as the best network in the analysis. Note that strictly tree-like topology (i.e., *h* = 0) performs poorly when compared to network topologies (i.e., *h >* 0) in general.

**Figure 5:**
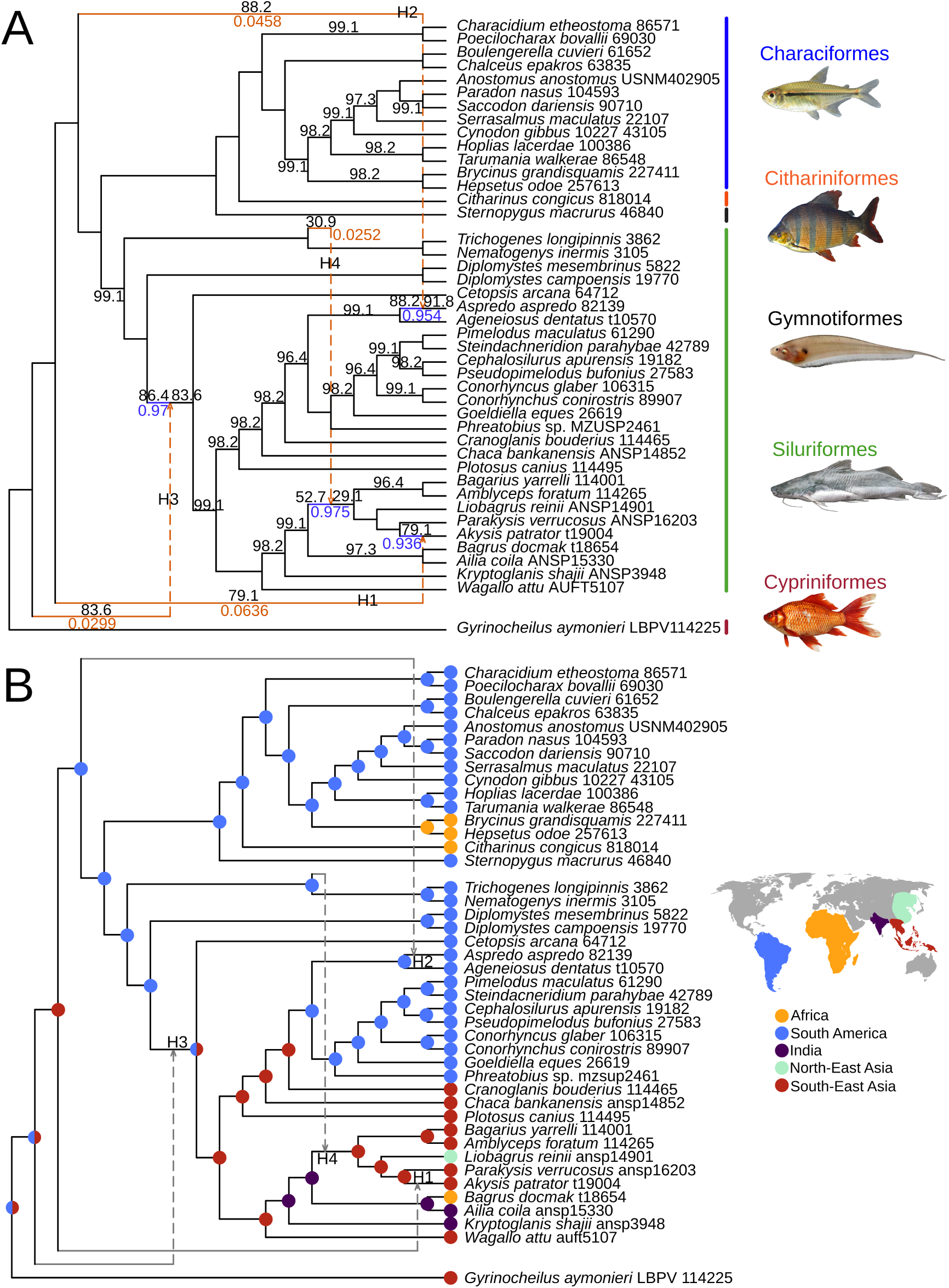
A. Phylogenetic network with four reticulations estimated using SNaQ. Support values on branches are bootstrap frequencies. Major hybrid edge on blue, minor hybrid edge in orange. Values are *γ* for the minor edge and 1 *− γ* for the major edge. B. Biogeographic reconstruction on the phylogenetic network using Fitch parsimony and five biogeographic regions. Internal nodes with more than one color represent ambiguous ancestral state reconstruction.

### 4.3 Uncertainty

Bootstrap support for the tree-like part of the chosen topology was relatively high, with values above 95 (Figure 5A). Bootstrap support for the parent and child nodes of a hybrid edge were variable. In some cases, the origin of the reticulation had high support (e.g., 83.6 in H3; Figure 5A), while in others it was the insertion with high support (e.g., 91.8 in H2; Figure 5A). In some other cases, the exact combination of origin and insertion of a reticulation had high support (e.g., 79.1 in H1; Figure 5A). Finally, H4 had overall low support in all features: origination, insertion, and exact combination (30.9, 52,7, and 29.1; Figure 5A).

Phylogenetic network uncertainty measured with the relative frequency of distinct topologies from the set of bootstrapped networks revealed that the best topology was present in 23% of the set, whereas other three topologies followed with 6.76% each (Figure 6). Focusing on a set of the four most frequent distinct topologies comprises nearly 50% of the cases, whereas preserving the best seven just raised the percentage to 55.4% (Supplementary Figure S1). These topologies differed in the presence of some hybrid edges, in line with the branch uncertainty bootstrap values; however, they did not differ in the tree-like part of the topology as measured by the Robinson-Foulds distance of their major trees, which was zero in all pairwise comparisons.

**Figure 6:**
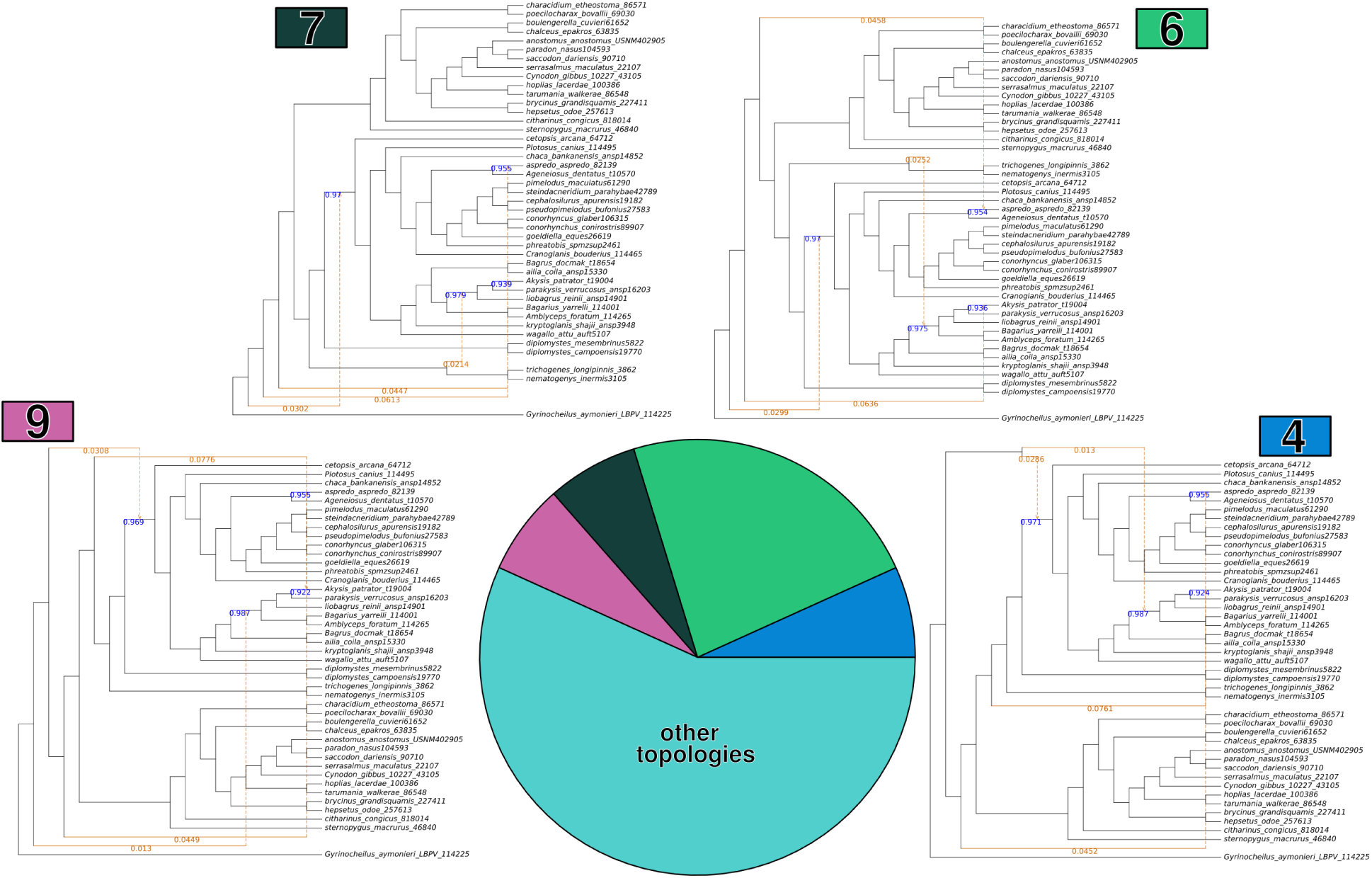
Topological uncertainty in phylogenetic network estimation measured with the topoFreq function of the tbea package. Topology 6 had the highest frequency with 23.0%, whereas topologies 4, 7, and 9 followed with 6.76% each. The remaining 56.76% represented other 31 topologies.

### 4.4 Sources of phylogenetic incongruence

We found that the species tree estimated by ASTRAL-4 was generally consistent with the major tree implied by the best network estimated by SNaQ (Figure 7). There were some discrepancies in the position of specific taxa, such as *Citharinus congicus*, *Phreatobius* sp., *Chaca bankensis*, and *Kryptoglanis shajii*. These taxa were consequential, as *Citharinus congicus* was the only representative of Cithariniformes in our taxon set. The other species were representatives of Siluriformes, for which interrelationships have been challenging to settle in the literature (Sullivan et al., 2006; de Pinna et al., 2025).

**Figure 7:**
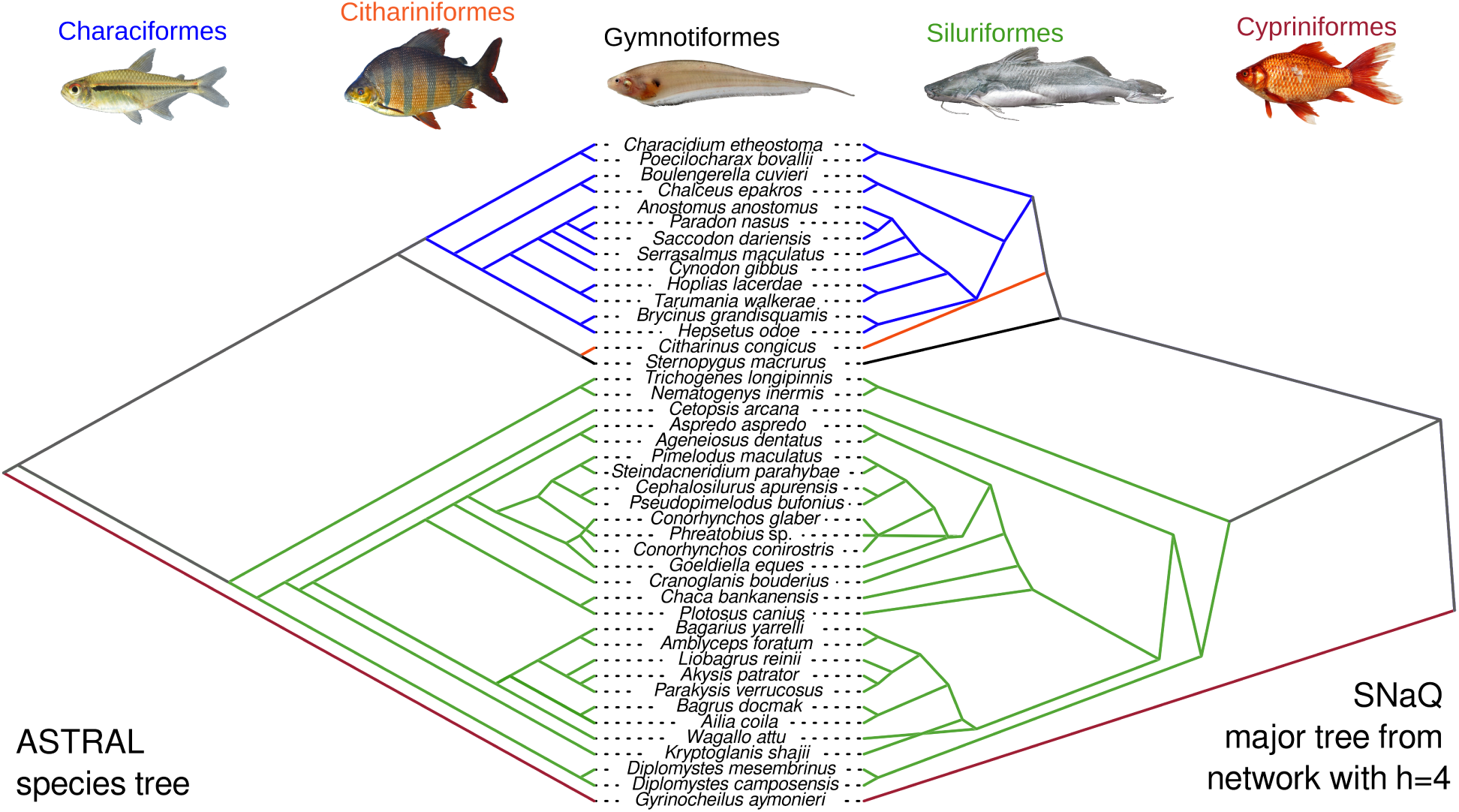
Cotangleplot between the ASTRAL-4 species tree under the coalescent, against the major tree from the phylogenetic network inferred with SNaQ.

Instances of moderate to high rates of incomplete lineage sorting (i.e., ILS-e *>* 30%) were found in 24 out of 40 branches, moderate instances of introgression/hybridization (i.e., IH-e *>* 5%) in one or two branches, or both cases in four nodes, following the thresholds in Shang et al. (2025) (Supplementary Table 2, Supplementary Figure 2). Also, branch support measured by local posterior probability was less than 100 in 15 out of 40 instances, and less than 90 in 13 (Figure 8). Although the exact branches in the species tree for which there was signal of reticulate evolution by IH-e may differ from branches where the phylogenetic network found hybrid edges, both approaches agreed on which regions of the ostariophysan phylogeny had a history of reticulate evolution (Figures 5,8).

**Figure 8:**
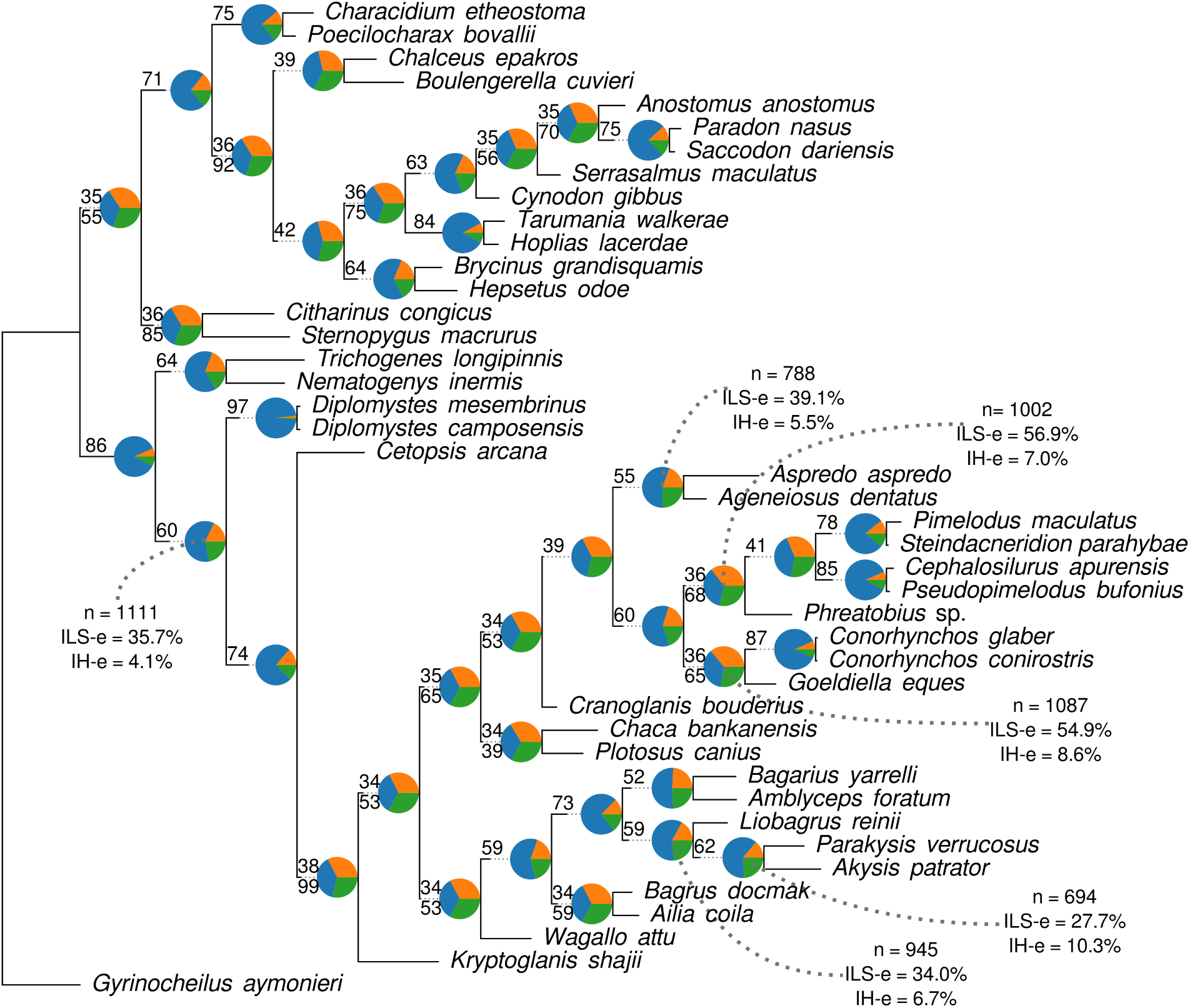
Species tree estimated with ASTRAL-4, summarizing sources of topological conflict with phytop. Pie charts are proportions of distinct quartet topologies q1, q2, and q3. The number above each branch is the proportion of q1, and the number below the local posterior probability (LPP). Only instances of LPP different from 100 are shown. Branches where positive signal of reticulate evolution was found show n = number of gene trees, ILS-e = proportion of gene trees that can be explained by ISL, IH-e = proportion of gene trees that can be explained by hybridization.

### 4.5 Biogeography

Biogeography was inferred on the best network with four hybrid edges (Figure 5B). Most internal nodes had their ancestral states reconstructed unambiguously. However, the root, node subtended by the root, and the suborder Siluroidei (defined as the most recent common ancestor of *Cetopsis arcana* and *Wallago attu*) had uncertainty in the reconstructed state, being either South America or South-East Asia. All reticulation events occurred between regions of the tree and terminals that share the same biogeographic area, except reticulation H4, which connects the suborder Loricarioidei and the clade defined by *Bagarius yarrelli* and *Akysis patrator* (approximately equivalent to the superfamily Bagroidea; de Pinna et al., 2025). The biogeographic model suggested two events between South America and Africa (the characiform family Alestidae and the order Cithariniformes), and one between India and Africa (*Bagarius docmak* and *Ailia coila*).

## 5 Discussion

The results from our analyses of reticulate evolution are partially consistent with previous findings from tree-based phylogenomic analyses of ostariophysan orders and highlighted reticulate evolution as a potential cause of past disagreements. Key differences from both the tree of blobs (Figure 3) and the ASTRAL-4 species tree (Figure 8) recovered Cithariniformes as sister to Gymnotiformes, with Characiformes and Siluriformes as successive groups, a novel relationship not found in previous studies (Figure 1) but most similar to Dai et al. (2018) with a swapped position of Characiformes and Cithariniformes. On the other hand, the major tree implied by the preferred phylogenetic network, that is, the network topology devoid of hybrid edges (Figures 5A,7), is similar to the works of or to Saitoh et al. (2003) and Orti & Meyer (1996) that recovered Siluriformes as sister to the clade (Gymnotiformes,(Characiformes + Cithariniformes)). Another finding here is the monophyly of Characiformes + Cithariniformes, consistent with the results of several previous studies (Saitoh et al., 2003; Lenglin et al., 2025; Fink & Fink, 1981; Arcila et al., 2017; Betancur-R et al., 2017; Betancur-R. et al., 2019; Hughes et al., 2018). Arcila et al. (2017) and Betancur-R. et al. (2019) devoted extensive effort to explore ways to better understand the interrelationships of ostariophysans in presence of conflicting phylogenomic signal, and provided useful methodological and empirical suggestions for balancing taxon and gene sampling, although both studies were based on the premise that a tree is the correct topology. Our results suggest that the presence of reticulate evolution may underlay variable conclusions reached by otherwise similar data and approaches, and offer an alternative view of using phylogenetic networks as a way forward for resolving recalcitrant groups in the fish tree of life.

It is important to note that ILS was found to be pervasive in the reduced dataset, and that signal of reticulate evolution from multiple sources (e.g., IH-e proportion assessed by phytop, the tree of blobs, and the composite likelihood score heuristics in SNaQ) explain why the ostariophysan interrelationships have been persistent issues in fish phylogenetics. First, the extensive gene–tree to species–tree discordance makes it challenging for methods such as maximum likelihood to estimate the correct species tree (Degnan & Rosenberg, 2009; Tabatabaee et al., 2023; Lanfear & Hahn, 2024; Zhang et al., 2025). Second, a strictly bifurcating phylogenetic tree was rejected for ostariophysans based on our data, and tree-based models can fail to recover the major tree topology group (Solís-Lemus & Ané, 2016; Kong et al., 2025), especially when ILS is high (Jiao et al., 2020). Because our SNaQ analyses account for ILS and reticulate evolution, we suggest the SNaQ ostariophysan hypothesis should be preferred over our ASTRAL analyses and similar phylogenomic hypotheses that did not account for reticulate evolution.

Phylogenetic network inference has long been identified as a computational problem more difficult than phylogenetic tree inference (Solís-Lemus & Ané, 2016; Wen et al., 2018). The approach employed in the present study with composite likelihood has historically involved successive point estimation and then bootstrapping, but this has the drawback of restricting the incorporation of uncertainty to the measurement of branch support. Our approach incorporated data uncertainty into network inference, providing a single step to network estimation and evaluating uncertainty among branch and competing network topologies. The use of topological frequency in a way similar to Bayesian phylogenetics is not typical, but we propose it is useful for assessing support among networks as full-likelihood Bayesian approaches for phylogenies even of moderate size is not feasible (Wen et al., 2018; Kong et al., 2025). This approach pairs well with heuristic methods such as SNaQ, and has some similarities to the multi-step heuristic searches under parsimony using TNT (Goloboff & Catalano, 2016).

As mentioned earlier in this article, a divergence time estimation analysis is key for analyzing networks using probabilistic models. Some alternatives exist, such as using least squares (Bastide et al., 2018) from genetic distance data, which work in datasets following a strict clock (Karimi et al., 2020). However, deep divergences and relaxed clocks do not allow us to find ultrametric topologies in some time unit instead of coalescent units or expected numbers of substitutions per site. Although Bayesian methods for divergence time estimation on phylogenetic network such as the SpeciesNetwork package (Zhang et al., 2018) or BPP4 (Flouri et al., 2020) can work for small datasets (Tiley et al., 2023), these methods were not possible for our data and relaxed clocks on networks have not been explored. Our alternative was to use Fitch parsimony and ancestral state reconstruction. Although not ideal, it is an entry point to biogeography on networks, and allows us to perform basic checks on whether reticulate evolution leads to some paradigm shifts in the evolution of major groups.

Several studies have tackled the biogeography of ostariophysans either discussing zoogeographic patterns (Briggs, 2005), or using probabilistic methods and phylogenetic trees (Saitoh et al., 2003; Chen et al., 2013; Lenglin et al., 2025; Liu et al., 2025; Brownstein et al., 2026; Hays et al., 2026). Several works support an origin of otophysans (i.e., Ostariophysi exclusive of Gonorhynchiformes) in Pangea (Saitoh et al., 2003; Chen et al., 2013; Lenglin et al., 2025), whereas others suggest a younger origin time for the group, challenging the role of the break up of Panegea in the diversification of the group (Liu et al., 2025; Brownstein et al., 2026). Because of the Afro-American distribution of Characiformes+Cithariniformes, some authors have suggested an important role of the Gondwanan break up in the diversification of this group (Saitoh et al., 2003). The cosmopolitan distribution of Siluriformes has also been suggested for the time of origin of Siluriformes as coeval with Pangea or at least Gondwana (Gayet & Meunier, 2003), and the Laurasian distribution of Cypriniformes as evidence of a Pangean origin of Otophysi (Chen et al., 2013). Here we find that the relationship between South American and West African taxa suggests two geodispersal events affecting the family Alestidae, and the order Cithariniformes, caused by the western Gondwana breakup producing South America and Africa. An event involving Africa and India is suggested to be related to the earlier relationship between the Indian plate and the African plate, where the former displaced away from the latter starting at around 100Ma and colliding to Asia in its modern configuration at around 55Ma. All of the previous biogeography studies have relied on phylogenetic trees for estimating the biogeography of ostariophysan fishes. It is still unclear to what extent their conclusions can differ once phylogenetic networks are used instead of phylogenetic trees. However, if such models behave like phylogenetic comparative methods, we can expect bias in estimation as has been found when the reticulate structure of topologies is ignored (Bastide et al., 2018; Morley et al., 2026).

It is expected that the approach proposed herein will help tackle large problems in phylogenetic network estimation while simultaneously accounting for uncertainty in data as well as topological and branch uncertainty. We showed that reticulate evolution can be important in understanding the interrelationships of ostariophysans, and suggest it explains phylogenetic conflicts in previous studies using tree inference methods where sampling of taxa or loci was not a large concern (Alda et al., 2019; Arcila et al., 2017; Arroyave et al., 2013; Betancur-R et al., 2017; Chakrabarty et al., 2017; Chen et al., 2013; Dai et al., 2018; Fink & Fink, 1981; Hughes et al., 2018; Rivera-Rivera & Montoya-Burgos, 2018). We anticipate that more detailed studies of reticulate evolution in ostariophysans, especially under more compelling models of trait evolution and biogeography, will provide a more complete picture of fish evolution.

## 6 Acknowledgments

This research was carried out using services provided to the Ballen Group by the Open Science Grid Consortium (Pordes et al., 2007; Sfiligoi et al., 2009; OSG, 2006, 2015), which is supported by the National Science Foundation awards #2030508 and #2323298. GAB was funded through a postdoctoral fellowship by FAPESP (#2023/07838-1) and a Thematic Project Grant by FAPESP awarded to C. de Oliveira (#2020/13433-6). We are thankful to Alongklod Tanomtong and Weerayuth Supiwong (Khon Kaen University), Armando Ortega-Lara (INCIVA), Bruno F. Morales (UFGD), Caleb McMahan, Susan Mochel, and Kevin Swegel (FMNH), Carlos Lucena (MCP), Carolina Doria (UFRO-ICT), Cecile Gama (IEPA), Francisco Villa-Navarro (CZUT-IC), Herńan Ĺopez-Ferńandez and Juan Albornoz-Garźon (UMMZ), Herńan Ortega (UNMSM), Jonathan W. Armbruster (AUM), Jorge García-Melo (Univ of Ibagué), Lucia Rapp Py-Daniel (INPA), Marcelo R. S. Melo (IO-USP), María Castillo (STRI), Mario de Pinna and Murilo N. Pastana (MZUSP), Mark H. Sabaj and Mariangeles Arce (ANSP), Mary Burridge (ROM), Mauro Nirchio (UT Machala), Nathan K. Lujan (ROM), Ramiro Barriga (MEPN), and Ryan Thoni and Radford Arrindell (AMNH) for curatorial assistance and loan of materials. We would also like to thank everyone who contributed to the DNA sample extraction and handling stages. Thanks to Erin Molloy, Cécile Ané, Hector Baños, Josh Justison, and Sandra Reinales for their help and feedback in computational phylogenetics. Thanks to the sources of the pictures used in the figures made available under free licenses over Wikimedia Commons: *Carassius*, *Chanos*, *Distichodus*, *Eigenmannia*, *Hyphessobrycon*, *Phractocephalus*, and Continents.

## 6.1 Code and data availability

All the phylogenomic analyses were run on the Open Science Grid Consortium and the Open Science Data Federation (Pordes et al., 2007; Sfiligoi et al., 2009; OSG, 2006, 2015). Biogeographic analysis and subsequent summarization of analyses was carried out on a laptop under Linux. Code and data are available on both Github (https://github.com/gaballench/ostariophysan_networks) and Zenodo (DOI 10.5281/zenodo.24090276). DNA alignments are available on Dryad (DOI XXXXX).

## Notes

### Competing Interest Statement

The authors have declared no competing interest.

